# Beta-adrenergic signaling and T-lymphocyte-produced catecholamines are necessary for interleukin 17A synthesis

**DOI:** 10.1101/2024.06.05.597633

**Authors:** Tatlock H. Lauten, Safwan K. Elkhatib, Tamara Natour, Emily C. Reed, Caroline N. Jojo, Adam J. Case

**Author notes:** **Corresponding author:** Adam J. Case, PhD, Associate Professor, Department of Psychiatry and Behavioral Sciences, Department of Medical Physiology, 8447 Riverside Pkwy, MREB2 3414, Bryan, TX 77807.

## Abstract

**Background:** Post-traumatic stress disorder (PTSD) is a debilitating psychological disorder that also presents with neuroimmune irregularities. Patients display elevated sympathetic tone and are at an increased risk of developing secondary autoimmune diseases. Previously, using a preclinical model of PTSD, we demonstrated that elimination of sympathetic signaling to T-lymphocytes specifically limited their ability to produce pro-inflammatory interleukin 17A (IL-17A); a cytokine implicated in the development of many autoimmune disorders. However, the mechanism linking sympathetic signaling to T-lymphocyte IL-17A production remained unclear.

**Methods:** Using a modified version of repeated social defeat stress (RSDS) that allows for both males and females, we assessed the impact of adrenergic receptor blockade (genetically and pharmacologically) and catecholamine depletion on T-lymphocyte IL-17A generation. Additionally, we explored the impact of adrenergic signaling and T-lymphocyte-produced catecholamines on both CD4+ and CD8+ T-lymphocytes polarized to IL-17A-producing phenotypes ex vivo.

**Results:** Only pharmacological inhibition of the beta 1 and 2 adrenergic receptors (β1/2) significantly decreased circulating IL-17A levels after RSDS, but did not impact other pro-inflammatory cytokines (e.g., IL-6, TNF-α, and IL-10). This finding was confirmed using RSDS with both global β1/2 receptor knock-out mice, as well as by adoptively transferring β1/2 knock-out T-lymphocytes into immunodeficient hosts. Furthermore, ex vivo polarized T-lymphocytes produced significantly less IL-17A with the blockade of β1/2 signaling, even in the absence of exogenous sympathetic neurotransmitter supplementation, which suggested T-lymphocyte-produced catecholamines may be involved in IL-17A production. Indeed, pharmacological depletion of catecholamines both in vivo and ex vivo abrogated T-lymphocyte IL-17A production demonstrating the importance of immune-generated neurotransmission in pro-inflammatory cytokine generation.

**Conclusions:** Our data depict a novel role for β1/2 adrenergic receptors and autologous catecholamine signaling during T-lymphocyte IL-17A production. These findings provide a new target for pharmacological therapy in both psychiatric and autoimmune diseases associated with IL-17A-related pathology.

## Introduction

The similarities between the nervous and immune systems are striking, with discoveries of new neuroimmune interactions being elucidated seemingly every day. It has become overwhelmingly appreciated that immune cells are able to respond to classical neurotransmitters, and conversely, neurons may be controlled by classical cytokines (Nance and Sanders, 2007; Padro and Sanders, 2014). Additionally, immune cells are able to produce classical neurotransmitters, while neurons are able to produce classical cytokines (Elkhatib and Case, 2019; Flierl et al., 2007; Olofsson et al., 2016; Pavlov and Tracey, 2005; Rosas-Ballina et al., 2011), thus, blurring the lines between what was once believed to be system-specific communication molecules. One significant early observation in neuroimmune communication was the finding that lymphoid organs appear to be innervated exclusively by sympathetic nerve fibers (Felten et al., 1987; Felten et al., 1985; Felten and Olschowka, 1987; Livnat et al., 1987), which suggested a potential role for catecholamine-mediated immune regulation. Indeed, numerous studies have demonstrated that beta-adrenergic signaling, primarily through the beta 2 (β2) adrenergic receptor, is able to control different types of immune cell trafficking, proliferation, and cytokine production (Kohm and Sanders, 2001; McAlees et al., 2011; Ramer-Quinn et al., 1997; Ramer-Quinn et al., 2000; Sanders, 2012; Simon et al., 2023). T-lymphocyte adrenergic signaling has also been specifically investigated, with many studies confirming the importance of the β2 receptor, but also demonstrating alternative regulation by other adrenergic receptors (Case et al., 2016; Deo et al., 2013; Huang et al., 2015; Takayanagi et al., 2012; Xu et al., 2019). However, few consensuses exist regarding the sympathetic regulation of T-lymphocytes given that the majority of studies have demonstrated that adrenergic responses are highly complex given the diversity of adrenergic receptors, specific types and concentration of catecholamines at play, and the unique (patho)physiological context that is being studied (Elkhatib and Case, 2019; Padro and Sanders, 2014).

One disease that highlights the importance of autonomic-immune interactions is post-traumatic stress disorder (PTSD), in which patients exhibit elevated sympathetic nervous system activity as well as systemic inflammation. Moreover, PTSD patients are at significantly elevated risks of developing comorbid autoimmune and cardiovascular diseases (Boscarino, 2004; Boscarino et al., 2010; Burg and Soufer, 2016; Dong et al., 2004; Mikuls et al., 2013), which may be due to the dysregulated neuroimmune interactions. Importantly, in a preclinical mouse model of PTSD (i.e., repeated social defeat stress; RSDS), we have recapitulated both the enhanced sympathoexcitation and inflammation seen in the human disease (Elkhatib et al., 2020; Moshfegh et al., 2019). Of note, interleukin 17A (IL-17A) is highly elevated in circulation in both mice and humans after psychological trauma, which is important given this proinflammatory cytokine’s role in both cardiovascular and autoimmune diseases (Bedoya et al., 2013; Karbach et al., 2014; Madhur et al., 2011; Madhur et al., 2010; Nguyen et al., 2013; von Vietinghoff and Ley, 2010; Zambrano-Zaragoza et al., 2014). We have further demonstrated that the spleen is a primary source of T-lymphocytes producing IL-17A after psychological trauma, and that targeted sympathetic denervation of this lymphoid organ is sufficient to attenuate IL-17A production during RSDS (Elkhatib et al., 2021). While this observation suggests an important role for neural-derived neurotransmission in RSDS-induced IL-17A synthesis, we have also found that T-lymphocytes lacking tyrosine hydroxylase (the rate limiting enzyme in catecholamine production) were limited in their ability to produce IL-17A after RSDS (Elkhatib et al., 2022). Together, this suggests that both neural and immune-derived adrenergic signaling are important in IL-17A production from T-lymphocytes, but the details involved in this regulation are unknown.

To explore this further, herein, we investigated the impact of adrenergic antagonism on RSDS-induced IL-17A. During this work, we repeatedly observed significant attenuation of in vivo systemic IL-17A with combined β1/2 blockade both with and without RSDS, but not with blockade of other adrenergic receptors. Moreover, β1/2 blockade blunted T-lymphocyte synthesis of IL-17A ex vivo during IL-17A-polarizing conditions, even in the absence of any exogenous catecholamine supplementation. These findings suggested that T-lymphocyte catecholamine production and autocrine/paracrine signaling are essential for the generation of IL-17A, and indeed, depletion of catecholamines both in vivo and ex vivo abrogated the ability of T-lymphocytes to specifically generate this proinflammatory cytokine. Overall, these data highlight a previously undescribed and critical role for β-adrenergic signaling in T-lymphocyte IL-17A production, which may have a significant clinical impact for IL-17A-associated diseases such as PTSD, autoimmunity, and cardiovascular pathologies.

## Materials and Methods

### Mice

Wild-type C57BL/6J (#000664; shorthand WT), β1/2 adrenergic receptor knock-out (#003810; shorthand β1/2^-/-^), Rag2 knock-out mice (#008449; shorthand Rag2^-/-^), and estrogen receptor 1 alpha cre (#017913; shorthand Esr1-cre) mice were obtained from Jackson Laboratories (Bar Harbor, ME, USA). CD1 mice were purchased from Charles River (#022, Wilmington, MA, USA). Esr1-cre mice were backcrossed to CD1 mice >10 generations to create a congenic CD1 background strain. All mice were bred in house to eliminate shipping stress and microbiome shifts, as well as co-housed with their littermates (≤5 mice per cage) prior to the start of experimentation to eliminate social isolation stress. Mice were housed with standard pine chip bedding, paper nesting material, and given access to standard chow (#8604 Teklad rodent diet, Inotiv, West Lafayette, IN, USA) and water ad libitum. Male and female experimental mice between the ages of 8-12 weeks were utilized in all experiments, but no sex differences were observed so data are presented as pooled independent of sex. Experimental mice were randomized, and when possible, experimenters were blinded to the respective cohorts until the completion of the study. Mice were sacrificed by pentobarbital overdose (150mg/kg, Fatal Plus, Vortech Pharmaceuticals, Dearborn, MI, USA) administered intraperitoneally. All mice were sacrificed between 7:00 and 9:00 Central Time to eliminate circadian rhythm effects on T-lymphocyte function. All procedures were reviewed and approved by Texas A&M University Institutional Animal Care and Use Committees.

### Adapted repeated social defeat stress (RSDS)

The standardized model of RSDS relies on the aggression of males (Golden et al., 2011), therefore, precludes the use of females. To address this limitation, we have recently adapted a novel model of RSDS that utilizes chemogenetically altered aggressive mice that indiscriminately show aggression to both male and female experimental mice (Takahashi et al., 2017). Briefly, Esr1-cre mice were stereotactically injected with activating Designer Receptors Exclusively Activated by Designer Drugs (DREADDs) viral constructs (pAAV-hSyn-DIO-hM3D(Gq)-mCherry (AAV2); #44361-AAV2, Addgene, Watertown, MA, USA) into the aggression center of the ventromedial hypothalamus (VMHvl) (AP, −1.5; ML, □ ± □ 0.7; DV, −5.7□mm from bregma). Four weeks post-injection, mice were screened for aggression after administration of clozapine-N-oxide (CNO; 1mg/kg delivered intraperitoneally). Mice that demonstrated indiscriminate aggression towards both male and female experimental mice were utilized in subsequent RSDS experiments. For RSDS, experimental mice placed in the home cage of an Esr1-cre mouse (pre-injected with CNO) to induce a physical confrontation and fear induction for 1 minute given the highly aggressive nature of the chemogenetically-modified Esr1-cre mice. Following the aggressive interactions, a clear perforated divider was placed into the cage and both mice remained co-housed for 24 hours. This process was repeated with a new aggressive Esr1-cre mouse daily for 5 consecutive days. Control mice were pair housed in the absence of an Esr1-cre mouse for the duration of the protocol. Mice were continuously monitored for visual wounding or lameness; None of the mice utilized herein met the threshold for exclusion (wounds >1cm or any lameness).

### In vivo treatment regimens

Doxazosin (#77883-43-3, Thermo Fisher Scientific, Houston, TX, USA) an α1 antagonist, propranolol (#4199-10-4, Sigma Aldrich, St. Louis, MO, USA), a β1/2 receptor antagonist, or SR59230A (#21407-25, Cayman Chemical, Ann Arbor, MI, USA), a β3 antagonist, were reconstituted in sterile isotonic (0.9%) saline solution and delivered by >14-day delivery subcutaneous osmotic minipumps (ALZET Osmotic Pumps, #1002-100uL, Cupertino, CA, USA) using previously established doses of 5 mg/kg (Baranski et al., 2013; Yi et al., 2011). Drug or control (i.e., saline only) minipumps were surgically implanted dorsally and subcutaneously in mice three days before the start of the RSDS protocol. Reserpine (Sigma-Aldrich, #83580-5G, St. Louis, MO, USA), a catecholamine depletion agent, was administered via intraperitoneal injection prior to RSDS protocol using a previously established dose (Arora and Chopra, 2013; Ishola et al., 2014). Control mice received an equal volume of vehicles (i.e., DMSO only) prior to RSDS protocol.

### T-Lymphocyte Immunophenotyping

Flow cytometric immunophenotyping was performed as previously described (Elkhatib et al., 2022). Briefly, spleens were harvested, physically dissociated into single cell suspensions, and erythrocyte depleted. Cells were incubated at 37°C for 4 h in RPMI media supplemented with phorbol 12-myristate-13-acetate (PMA; 10 ng/mL), ionomycin (0.5 mg/mL), and BD GolgiPlug Protein Transport Inhibitor (containing brefeldin A; 1 mg/mL; BD Biosciences, Franklin Lakes, NJ, USA). Following this, cells were washed and resuspended in PBS containing an amine-reactive viability dye (Live/Dead Fixable Cell Stain, #L34960, Thermo Fisher Scientific, Houston, TX, USA) for 30 min at 4°C. Antibodies targeting extracellular markers were then added prior to fixation/permeabilization (#00-5523-00, eBioscience, San Diego, CA, USA) and addition of antibodies targeting intracellular markers. Specific antibodies utilized: CD3ε PE-Cy7 (#25-0031-82, Thermo Fisher Scientific, Houston, TX, USA), CD8 Alexa Fluor 488 (#69-0041-82, Thermo Fisher Scientific, Houston, TX, USA), CD4 eFluor 506 (#69004182, Thermo Fisher Scientific, Houston, TX, USA), and IL-17A BV711 (#407-7177-82, Thermo Fisher Scientific, Houston, TX, USA). Cells were assessed using a 4-laser Attune NxT flow cytometer (Thermo Fisher Scientific, Houston, TX, USA), while data was analyzed using FlowJo analysis software (BD Biosciences, Franklin Lakes, NJ, USA).

### T-lymphocyte isolation and activation

T-lymphocytes from spleens were isolated using negative selection, as previously described (Moshfegh et al., 2019). Briefly, splenocytes in a single cell suspension were negatively selected for CD4+ or CD8+ T-lymphocytes using the EasySep Mouse CD4+ T-cell Isolation Kit or CD8+ T-cell Isolation Kit (StemCell Technologies #19852, #19853, Vancouver, BC, USA), respectively. Cell viability was measured via a Bio-Rad TC20 Automated Cell Counter using trypan blue exclusion, and cell purity was assessed via flow cytometry (average yield is approximately 5 x 10^6 cells/mouse and purity achieved is >90%). T-lymphocytes were cultured in RPMI media supplemented with 10% Fetal Bovine Serum, 10 mM HEPES, 2 mM Glutamax, 100 U/mL penicillin/streptomycin, and 50 µM of 2-mercaptoethanol.

### IL-17A-producing polarization

CD4+ and CD8+ splenic T-lymphocytes were polarized to IL-17A-producing phenotypes ex vivo by use of CellXVivo Mouse T_H_17 Cell Differentiation Kit (#CDK017, R&D Systems, Minneapolis, MN, USA). Briefly, CD4+ and CD8+ T-lymphocytes were isolated and cultured as described above with RPMI media supplemented with Dynabeads Mouse T-Activator CD3/CD28 (Gibco #11452D) at a ratio of 1:1 cell to beads, mouse IL-6 (#130-096-682, Miltenyi Biotech, Bergisch Gladbach, North Rhine-Westphalia, Germany), mouse IL-23 (#130-096-676, Miltenyi Biotech, Bergisch Gladbach, North Rhine-Westphalia, Germany), mouse IL-1β (#130-101-681, Miltenyi Biotech, Bergisch Gladbach, North Rhine-Westphalia, Germany), human TGF-β1 (#130-095-067, Miltenyi Biotech, Bergisch Gladbach, North Rhine-Westphalia, Germany), anti-mouse IL-4 antibody (#130-095-709, Miltenyi Biotech, Bergisch Gladbach, North Rhine-Westphalia, Germany) anti-mouse IFN-γ antibody (#130-095-729, Miltenyi Biotech, Bergisch Gladbach, North Rhine-Westphalia, Germany), and anti-mouse IL-2 antibody (#130-095-736, Miltenyi Biotech, Bergisch Gladbach, North Rhine-Westphalia, Germany). While these polarizing conditions were optimized for CD4+ T-lymphocytes, we have found that the same conditions significantly and sufficiently polarize CD8+ T-lymphocytes as well. Following 72 hours of culture and activation, T-lymphocytes were harvested for analysis.

### In vitro treatment regimens

Cell culture wells were supplemented daily with L-Norepinephrine (#AAL0808703, Thermo Scientific Chemicals, Houston, TX, USA), Propranolol (Sigma Aldrich, # 4199-10-4, St. Louis, MO, USA), Doxazosin (#77883-43-3, Thermo Fisher Scientific, Houston, TX, USA), and/or Atipamezole (Cayman Chemical, #9001181-25 Ann Arbor, MI, USA) reconstituted in sterile PBS. Concentrations used are denoted in specific figures and were based on previously published dose curves (Case et al., 2016).

### Catecholamine quantification

Dopamine, norepinephrine, and epinephrine levels in splenic lysate, plasma, and media were assessed using the 3-CAT research ELISA (Rocky Mountain Diagnostics, BAE-5600, Colorado Springs, CO, USA). All assays were completed according to the manufacturer’s protocol, with splenic lysate and media catecholamine concentration normalized to wet tissue weight and cell counts, respectively.

### cAMP quantification

Quantification of cAMP levels within cell lysates was assessed using the cAMP research ELISA (#ab290713, Cambridge, United Kingdom). All assays were completed according to the manufacturer’s protocol, with cell lysate normalized to live cell counts.

### Cytokine Analysis

Cytokines in plasma and media (normalized to cell counts) were measured as previously described (Elkhatib et al., 2020). Briefly, concentrations were assessed using custom Mesoscale Discovery U-Plex kits (Mesoscale Discovery, Rockville, MD, USA) per manufacturer’s instructions. Analysis was performed using a Mesoscale QuickPlex SQ 120 and analyzed using Mesoscale Discovery software.

### Statistical Analysis

A total of 107 animals (53 WT, 54 β1/2^-/-^) were utilized for the studies described herein. All data are presented as mean ± standard error of the mean (SEM) with sample numbers displayed as individual data points and N values are included within figure legends where appropriate. Data were first assessed for normality utilizing the Shapiro-Wilk test. Following this, data were analyzed using Mann-Whitney U-test, 1-way ANOVA, and 2-way ANOVA where appropriate (specific tests identified in the figure legend). All statistics were calculated using GraphPad Prism (V10, GraphPad).

## Results

### Inhibition of T-lymphocyte β-adrenergic signaling specifically impairs IL-17A expression

To investigate the effects of adrenergic signaling on RSDS-induced inflammation, surgically implanted osmotic minipumps delivering doxazocin (α1 blockade), propranolol (β1/2 blockade), or SR59230A (β3 blockade) were utilized throughout the duration of the RSDS paradigm. Only propranolol-infused animals demonstrated significant attenuation of circulating levels of IL-17A (**Figure 1A**) and IL-22 (data not shown) independent of RSDS. This decrease in IL-17A and IL-22 was highly specific, as other prominent systemic inflammatory cytokines reported elevated in RSDS (Elkhatib et al., 2020) remained unchanged or even elevated (**Supplemental Figure 1A-C**). Interestingly, β3 inhibition appeared to increase IL-17A at baseline to levels normally observed with RSDS, but no additional elevation was observed with RSDS (**Figure 1A**). To eliminate possible non-β-adrenergic or off-target effects of propranolol, we next utilized mice lacking β1 and β2 adrenergic receptors (i.e., β1/2^-/-^). After RSDS induction, circulating IL-17A levels significantly increased in WT animals as expected, however, β1/2^-/-^ animals showed minimal circulating IL-17A with or without RSDS (**Figure 1B**). Again, this effect was highly specific to IL-17A as other inflammatory cytokines were unaffected by the loss of β1 and β2 adrenergic receptors (**Supplemental Figure 1D-F**). To further investigate the role of β1/2 signaling directly on T-lymphocytes, a primary source of IL-17A, adoptive transfer of splenic T-lymphocytes from WT and β1/2^-/-^ donor animals was performed into T-lymphocyte-deficient Rag2^-/-^ host animals. Once again, Rag2^-/-^ mice receiving WT T-lymphocytes demonstrated the characteristic increase in circulating IL-17A after RSDS, but this effect was completely abolished in Rag2^-/-^ hosts receiving β1/2^-/-^ T-lymphocytes independent of RSDS (**Figure 1C**). Together, these data put forth strong evidence that β1/2-adrenergic signaling on T-lymphocytes is necessary for IL-17A expression.

**Figure 1.**
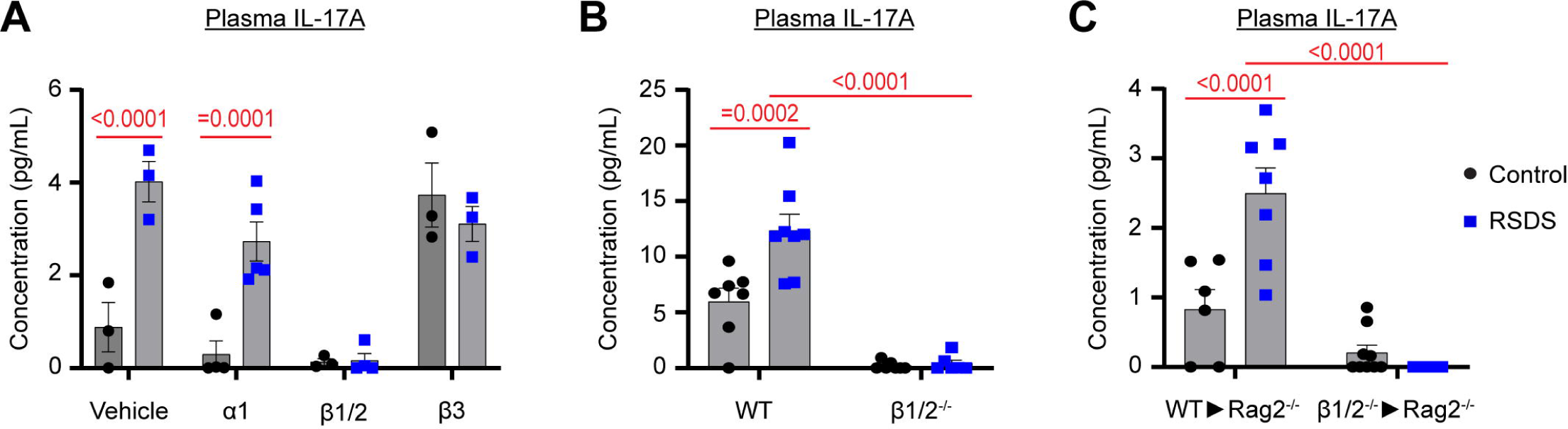
β1/2-blockade attenuates RSDS-induced circulating IL-17A. RSDS was performed in the presence/absence of either pharmacological or genetic adrenergic blockade followed by assessment of plasma IL-17A by Meso Scale Discovery assay. **A**. Pharmacological adrenergic blockade using doxazosin (α1), propranolol (β1/2), or SR59230A (β3). **B**. Global genetic β1/2 knockout (β1/2^-/-^). **C**. Adoptive transfer of β1/2^-/-^ T-lymphocytes into immunodeficient Rag2^-/-^ mice. Statistics by 2-way ANOVA with Bonferroni post-hoc.

After observing the effects of β1/2-adrenergic blockade on T-lymphocytes in vivo, we next examined the direct effects of β1/2 antagonism ex vivo. Splenic CD4+ and CD8+ T-lymphocytes were isolated from WT animals and cultured under IL-17A polarizing conditions in the presence or absence of propranolol. Propranolol treatment elicited significant decreases in IL-17A intracellular and secreted protein in both CD4+ and CD8+ T-lymphocytes compared to vehicle control (**Figure 2A-B, E-F**). The identical IL-17A inhibition was observed in both CD4+ and CD8+ T-lymphocytes from β1/2^-/-^ animals compared to WT T-lymphocytes (**Figure 2C-D, G-H**). These data suggest that β-adrenergic signaling on T-lymphocytes is essential for production of IL-17A, even in the absence of external sources of catecholaminergic neurotransmission (i.e., neurons).

**Figure 2.**
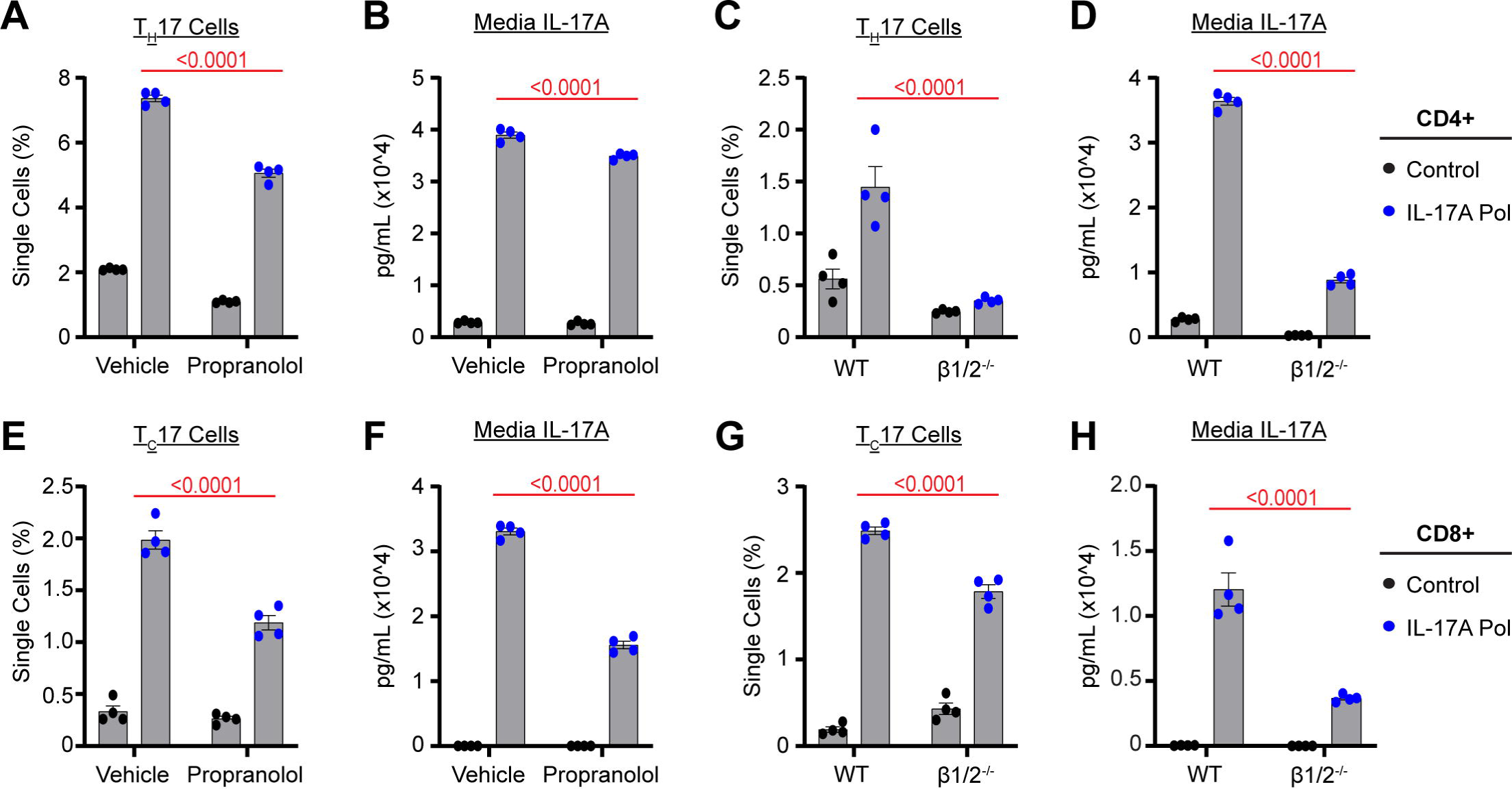
β1/2-blockade limits IL-17A production under ex vivo polarizing conditions. CD4+ or CD8+ T-lymphocytes were polarized under IL-17A-producing conditions ex vivo in the presence/absence of either pharmacological or genetic adrenergic β1/2 blockade followed by assessment of IL-17A+ cells by flow cytometry or secreted IL-17A by Meso Scale Discovery assay. **A-B**. CD4+ T-lymphocytes ± propranolol. **C-D**. β1/2^-/-^ CD4+ T-lymphocytes. **E-F**. CD8+ T-lymphocytes ± propranolol. **G-H**. β1/2^-/-^ CD8+ T-lymphocytes. Statistics by 2-way ANOVA with Bonferroni post-hoc.

### T-lymphocyte produced catecholamines are necessary for IL-17A

To further investigate the role of catecholamines in IL-17A synthesis, we treated animals with reserpine to cause a complete depletion of these neurotransmitters. As expected, reserpine treatment led to complete abrogation of catecholamines in the spleen (**Supplemental Figure 2A-C**). Furthermore, reserpine treatment led to significant attenuation of circulating IL-17A after RSDS compared to vehicle treated animals (**Supplemental Figure 2D**). While this data further confirms the necessity of catecholamines in IL-17A production, it does not provide evidence as to the source of the neurotransmitters needed to regulate this inflammatory cytokine.

Understanding that systemic reserpine treatment would impact both neuronal and immune cell catecholamine release, we next assessed the role of reserpine on T-lymphocytes in culture to further investigate the direct effects of catecholamine depletion from these cells. Strikingly, both CD4+ and CD8+ cells treated with reserpine under IL-17A polarizing conditions showed significant reduction in both IL-17A intracellular and secreted protein (**Figure 3A-D**), supporting the notion that T-lymphocyte produced catecholamines are essential for IL-17A production. To further explore possible intracellular signals linking catecholamine signaling to IL-17A production, cultured T-lymphocytes were treated with various adrenergic antagonists in combination with norepinephrine to assess intracellular cyclic AMP (cAMP) levels. Propranolol was the only antagonist that led to complete nullification of NE-driven intracellular cAMP in T-lymphocytes (**Figure 4A**). Similarly, β1/2^-/-^ T-lymphocytes possessed no appreciable increase in cAMP when treated with NE (**Figure 4B**), suggesting cAMP may be a critical second messenger in the generation of IL-17A. Indeed, T-lymphocytes cultured in IL-17A polarizing conditions demonstrated elevated intracellular cAMP compared to non-polarizing activating conditions, and propranolol was able to significantly eliminate this increase (**Figure 4C**). In summary, these data suggest that T-lymphocyte-produced catecholamines signaling through β1/2 adrenergic receptors and cAMP are necessary to modulate IL-17A generation in these adaptive immune cells.

**Figure 3.**
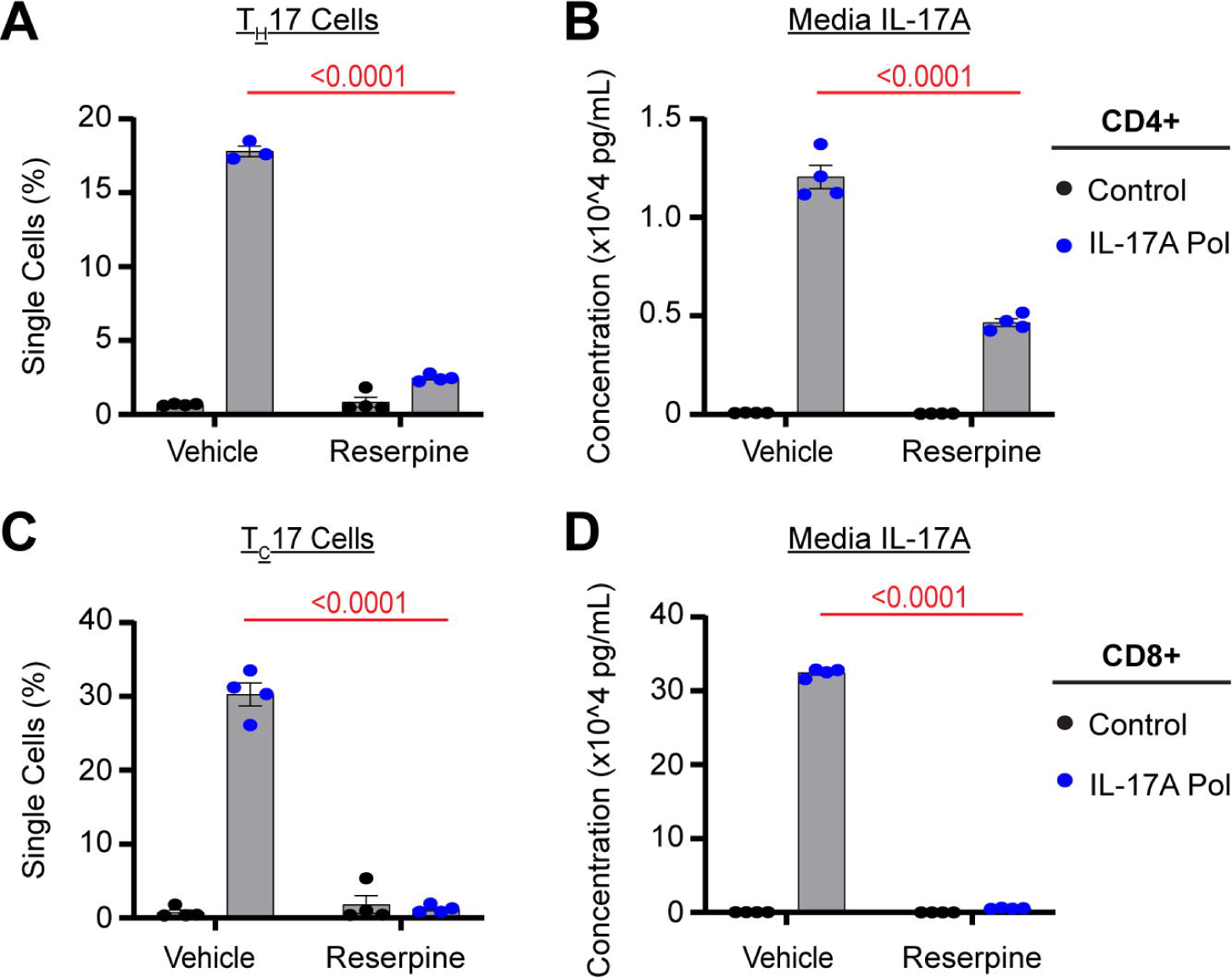
T-lymphocyte catecholamine depletion abrogates IL-17A production. CD4+ or CD8+ T-lymphocytes were polarized under IL-17A-producing conditions ex vivo in the presence/absence reserpine followed by assessment of IL-17A+ cells by flow cytometry or secreted IL-17A by Meso Scale Discovery assay. **A-B**. CD4+ T-lymphocytes ± reserpine. **C-D**. CD8+ T-lymphocytes ± reserpine. Statistics by 2-way ANOVA with Bonferroni post-hoc.

**Figure 4.**
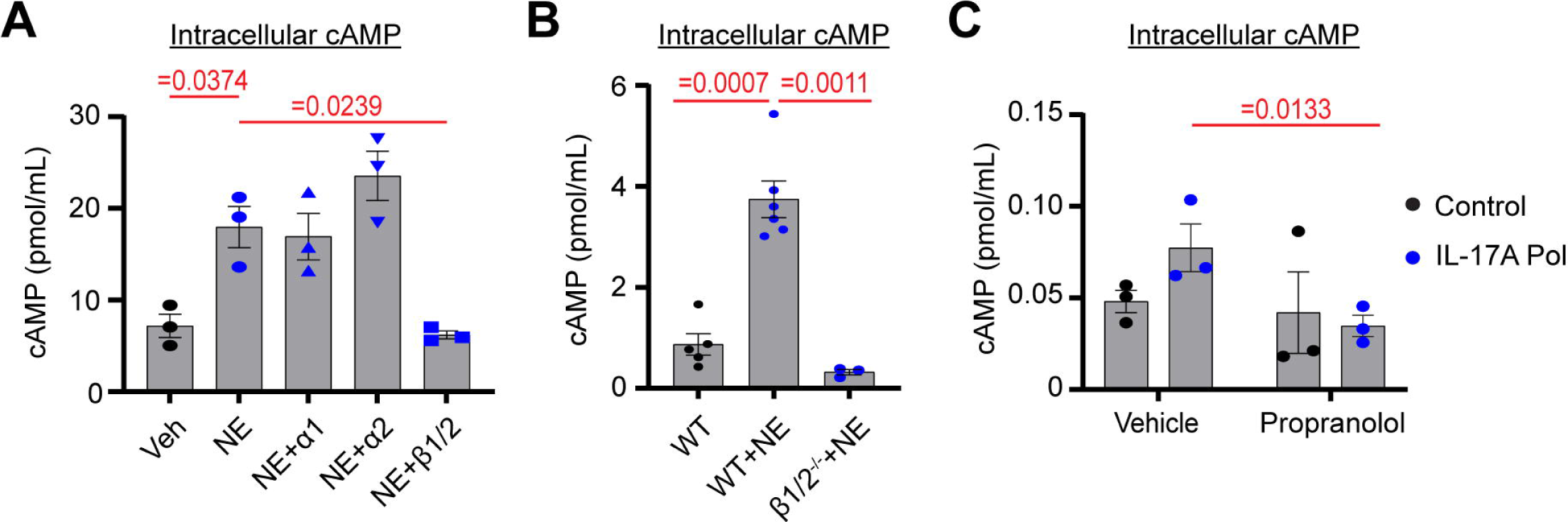
cAMP serves as a critical second messenger under IL-17A-producing conditions in T-lymphocytes. T-lymphocytes were cultured for one hour in the presence/absence of IL-17A polarizing conditions as well as exogenous NE and pharmacological/genetic blockade followed by assessment of intracellular cAMP. **A**. NE (1 µM) plus pharmacological α1 (doxazosin; 1 µM), α2 (atipamezole; 1 µM), or β1/2 (propranolol; 1 µM) blockade. **B**. NE plus β1/2 genetic knock-out. **C**. IL-17A polarizing conditions plus propranolol (1 µM). Statistics by 1-way ANOVA with Tukey post-hoc (**A-B**) or 2-way ANOVA with Bonferroni post-hoc (**C**).

## Discussion

In our previous investigations, we unveiled a pivotal connection between neural-derived catecholamines and T-lymphocyte generated IL-17A (Elkhatib et al., 2021). In attempts to better understand connections between psychological trauma and peripheral inflammation, we performed targeted splenic denervation in the context of RSDS. These studies elucidated that the absence of the splenic nerve led to a significant reduction of RSDS-induced IL-17A production by T-lymphocytes in this lymphoid organ. However, even in the absence of sympathetic innervation, the spleen still possessed detectable levels of catecholamines (Elkhatib et al., 2021), which suggested an alternative source of these neurotransmitters. Interestingly, previous work from our lab discovered a remarkably significant and positive association between tyrosine hydroxylase and IL-17A expression in T-lymphocytes (Moshfegh et al., 2019). Given this tight correlation and understanding that T-lymphocytes are able to generate their own neurotransmitters (Elkhatib and Case, 2019), we previously engineered a conditional genetic knockout of tyrosine hydroxylase specifically in T-lymphocytes. While the phenotype in these mice was relatively mild, the primary outcome of these studies demonstrated that T-lymphocytes lacking tyrosine hydroxylase, and thus the ability to make catecholamines, were less efficient in producing IL-17A compared to wild-type cells (Elkhatib et al., 2022). Together, these previous data put forth a model that IL-17A polarization is dictated by both neural and immune-derived catecholamines, but the details surrounding this regulation remained unclear.

It has been known for several decades now that T-lymphocyte function and differentiation may be controlled by β-adrenergic receptors, however, regulation by these autonomic receptors has proven highly dynamic and context dependent (Bellinger and Lorton, 2014; Sanders, 2012). Moreover, conflicting reports as to which β-adrenergic receptor regulates T-lymphocytes are pervasive in the literature. For example, it has been noted that T-lymphocytes predominantly express the β2 receptor (Sanders, 2012), therefore, most investigations in T-lymphocytes have focused on this receptor when examining T-lymphocyte regulation. β2 receptor mediated function in T-lymphocytes has been linked to proliferation (Guereschi et al., 2013; Madden et al., 1994; Maisel and Michel, 1990), cytokine production (Ramer-Quinn et al., 2000; Sanders et al., 1997), and polarization (Guereschi et al., 2013; Vida et al., 2011). In contrast, β1 receptor signaling has limited evidence of expression, but has been demonstrated to have functional relevance to T-lymphocytes as well, albeit less reported than β2 (Freier et al., 2010; Takayanagi et al., 2012) and seemingly more effective to the T-regulatory subtypes (Cosentino et al., 2007). For example, NE binding to the β1 receptor on T-lymphocytes exerted a suppressive effect on IFN-γ and TNF-α production (Takayanagi et al., 2012). Additionally, the β3 receptor has also been shown to regulate various aspects of immunity (Courties et al., 2015; Heidt et al., 2014), however, signaling via this receptor has not been explored specifically in T-lymphocytes.

A first possible explanation for these conflicting results may be due to the regulation of the specific receptors. Notably, polarization of T-lymphocytes to specific subtypes has been observed to alter the expression level of β receptors, particularly β2 (Sanders, 2012). Depending on the context of the specific study, the receptor density and ratio of β1 to β2 may dramatically alter the physiological outcome and explain why one receptor appears more dominant than the other in various situations. Second, both the β1 and β2 receptor signal through cAMP as a secondary messenger, which may suggest β1 and β2 receptors provide redundant or additive signaling on T-lymphocytes that allows for compensation in the event of loss of one receptor. Last, given that the β1 and β2 receptors possess different affinities for catecholamines, the specific neurotransmitter in question may play a significant role in determining which receptor predominates signaling in T-lymphocytes.

One area where there exists a notable paucity of literature is that of the impact of β receptors on T-lymphocytes that produce IL-17A. This cytokine has garnered significant attention given its connection to numerous autoimmune and cardiovascular diseases (Bedoya et al., 2013; Guzik et al., 2007; Herrera et al., 2006; Shao et al., 2018), as well as its success as a pharmacological clinical target (Blair, 2019, 2021; Reszke and Szepietowski, 2017). During our studies described herein, we struggled to identify any literature that had investigated links between β adrenergic receptors and IL-17A. However, one group has recently published a series of papers demonstrating the β2-agonist, terbutaline, could promote T_H_17 polarization in CD4^+^ T-lymphocytes from human peripheral blood mononuclear cells (Carvajal Gonczi et al., 2023; Carvajal Gonczi et al., 2017). These findings agree with our data presented here suggesting β receptors are necessary for complete IL-17A production. However, while the investigators were sufficiently able to utilize β2 stimulation to produce more IL-17A, our preliminary studies using specific β1 or β2 inhibitors failed to show a similar reduction in IL-17A compared to combined β1/2 blockade (data not shown). As previously hypothesized, this may be due to the relative level of cAMP generated by each respective receptor, and the blockade of only one may be compensated for by the presence of the other. This question, as well as many others in this emerging area of neuroimmunology, clearly warrants further exploration.

While our study provides compelling evidence supporting the role of β adrenergic receptors in the regulation of IL-17A, the specific molecular signals and pathways involved in this process remain somewhat elusive. By employing reserpine to deplete catecholamines in vivo, we effectively induced a state of IL-17A deficiency following RSDS, thus reinforcing the significance of these neurotransmitters. More importantly, through direct treatment of T-lymphocytes with reserpine, we were able to elucidate the importance of T-lymphocyte-generated catecholamines on IL-17A regulation. Our data unequivocally demonstrates a negative impact on IL-17A production in the absence of T-lymphocyte catecholamine release, consistent with findings from previous studies utilizing T-lymphocyte-specific knockouts of tyrosine hydroxylase (Elkhatib et al., 2022). However, the specific catecholamine(s) involved in these processes is currently unknown. The β receptors possess differential affinities for each of the catecholamines, but all are able to bind these receptors in some capacity. In our previous study using tyrosine hydroxylase knock-out T-lymphocytes, we were able to fully rescue the IL-17A deficit by exogenous addition of norepinephrine, while dopamine and epinephrine only provided partial rescue (Elkhatib et al., 2022). While compelling that norepinephrine is sufficient to restore the defect in IL-17A production, it does necessarily mean it is the neurotransmitter being produced by T-lymphocytes themselves. Measuring catecholamines produced from T-lymphocytes has posed a significant technical challenge given that they produce these neurotransmitters at much lower levels than neurons. This is an ongoing area of research in our laboratory, and we are currently in the process of developing novel techniques specifically for this purpose.

The reciprocal relationship between T_H_17 and Treg cells has been extensively investigated, with accumulating evidence proposing that their balance may underline the pathogenic mechanisms implicated in autoimmune diseases (Lee, 2018). Intriguingly, our administration of β1/2 antagonists (or genetic knock-outs) resulted not only in the lowering of IL-17A-positive cells as shown, but also a noticeable positive trend in the number of Foxp3+ polarized cells (data not shown), a classic marker of the Treg population (Kanamori et al., 2016). Notably, both T_H_17 and Treg cells share a common signaling pathway mediated by transforming growth factor beta (TGF-β), but it is the molecular context in which TGF-β is able to promote polarization to one type or the other (Zhu and Paul, 2010). For instance, TGF-β works in combination with IL-6 and IL-21 to promote IL-23 receptor expression, thereby favoring T_H_17 differentiation (Lee, 2018; Omenetti and Pizarro, 2015). Conversely, in the absence of IL-6 and IL-21, TGF-β suppresses IL-23R expression and promotes the development of Foxp3+ Tregs (Lee, 2018; Omenetti and Pizarro, 2015). Yet, even with these pathways becoming more understood, there is a lack of knowledge as to how β adrenergic receptors contribute to this polarization duality. Based on our findings presented herein, we postulate that intracellular cAMP levels may also be an additional critical component that contributes to the polarization context of TGF-β, but further exploration is currently ongoing to uncover how catecholaminergic signaling modulates the T_H_17/Treg balance.

The inhibition of IL-17A by way of β adrenergic pharmacological blockade is highly provocative given its potential translatability to IL-17A-related disorders (Berry et al., 2022; McGeachy et al., 2019). For instance, while beta blockers are utilized in an array of cardiovascular disorders due to their direct cardiac effects, it has yet to be examined if these same drugs impact levels of IL-17A in these patients. This is of significant importance given that IL-17A is often elevated with cardiovascular disease and may serve as an additional target for therapy (Mikolajczyk and Guzik, 2019; Rodrigues-Diez et al., 2021). Outside of cardiovascular diseases, IL-17A has been even stronger linked to the pathogenesis of various autoimmune diseases (Maddur et al., 2012; McGeachy et al., 2019). Secukinumab, a monoclonal inhibitory antibody to IL-17A, is already in clinical use for various autoimmune diseases (Blair, 2019, 2021; Reszke and Szepietowski, 2017). While this therapeutic approach has shown significant promise, its molecular target is that of fully formed IL-17A protein. In other words, if the source of the IL-17A is in continual production, then repeated administration of Secukinumab is necessary to limit circulating levels. The addition of beta blockers in these patients could serve as an adjuvant therapy to restrain the production of IL-17A, which would limit the effects of the cytokine by two therapeutic approaches. However, consideration must be given to potential adverse effects associated with beta blockade therapy, particularly when utilizing agents targeting both β1 and β2 adrenergic receptors. For example, β2-AR blockade alone can lead to various health issues, often relating to the limiting of bronchodilation in the pulmonary system (Billington et al., 2017). Additionally, the differential effects of β1 versus β2 blockade on IL-17A production have not been fully elucidated. It is plausible that β1 blockade alone may not yield the same blunting effect on IL-17A production observed with combined β1 and β2-AR blockade, as the regulation of intracellular signaling pathways, such as cAMP, may not significantly impact cytokine production. Furthermore, our observations regarding β3 blockade suggest that the removal of a singular receptor may exacerbate IL-17A cytokine production, underscoring the complex interplay between β adrenergic receptors and immune modulation.

In conclusion, this study sheds light on novel neuroimmune mechanisms governing IL-17A production. These findings raise pertinent questions regarding the broader therapeutic effects of currently utilized beta blockers beyond their known therapeutic actions. While questions still remain regarding the complete mechanism of action of how β receptors modulate IL-17A, it does not diminish enthusiasm for the potential of these in clinical therapy for IL-17A-related disorders.

## Acknowledgments

This work was supported by the National Institutes of Health (NIH) R01HL158521 (AJC). This publication’s contents are the sole responsibility of the authors and do not necessarily represent the official views of the NIH.

## Author Contributions

THL, SKE, and AJC designed research studies. All authors conducted experiments, acquired data, and/or performed analyses. THL and AJC wrote the manuscript. All authors approved final version of the manuscript. AJC provided funding and experimental oversight.

