## Supplemental Data for "Beta-adrenergic signaling and T-lymphocyte-produced catecholamines are necessary for interleukin 17A synthesis"

**Title**

Tatlock H. Lauten^1,2^, Safwan K. Elkhatib^3^, Tamara Natour^1,2^, Emily C. Reed^1,2^, Caroline N. Jojo^1,2^, and Adam J. Case^1,2*^

**Affiliations:**

^1^Department of Psychiatry and Behavioral Sciences, Texas A&M University, Bryan, TX, United States

^2^Department of Medical Physiology, Texas A&M University, Bryan, TX, United States

^3^Department of Anesthesiology, Perioperative, and Pain Medicine, Brigham and Women's Hospital, Boston, MA

***Corresponding author:**	Adam J. Case, PhD

Associate Professor

Department of Psychiatry and Behavioral Sciences

Department of Medical Physiology

8447 Riverside Pkwy

MREB2 3414

Bryan, TX 77807

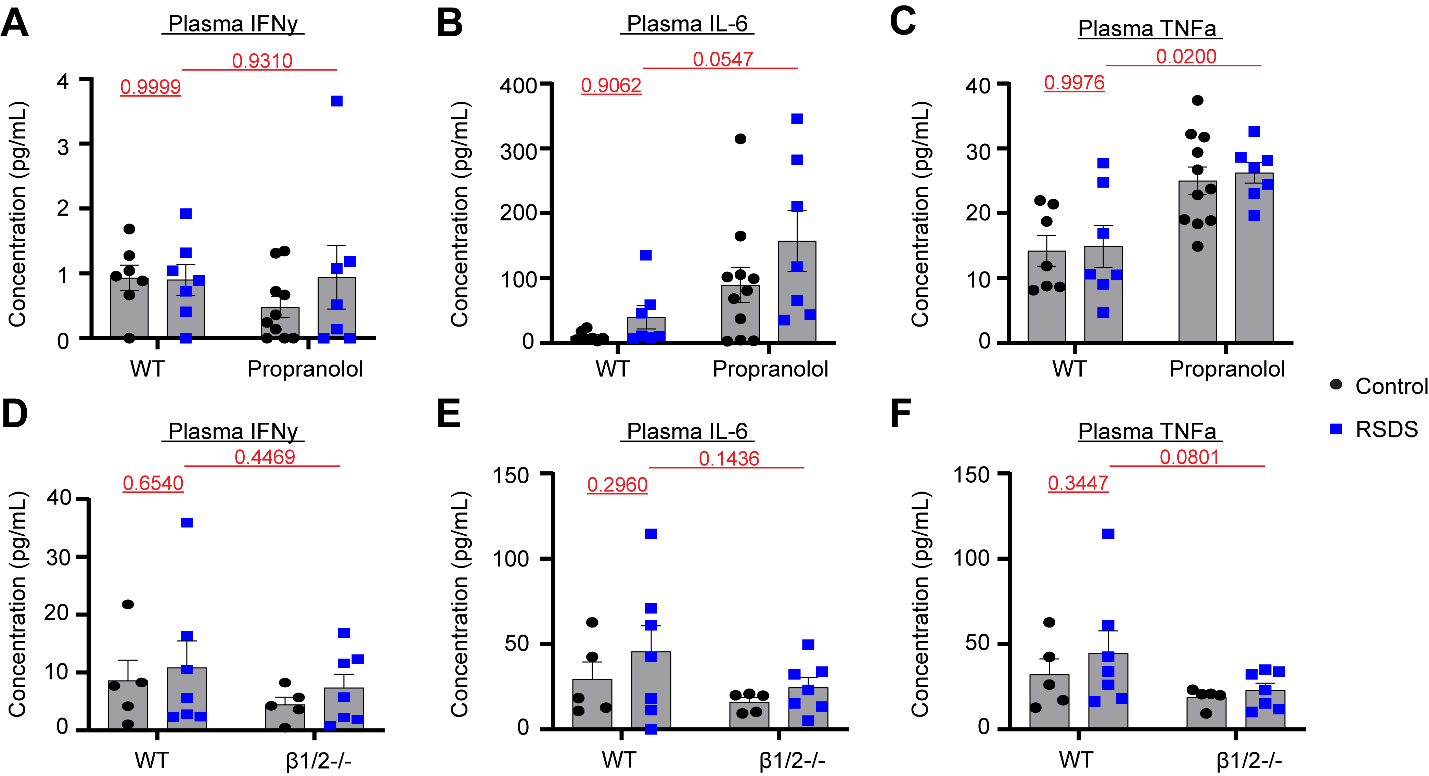

**Supplemental Figure 1. β1/2-blockade does not impact other RSDS-induced circulating cytokines.** RSDS was performed in the presence/absence of either pharmacological or genetic β1/2 blockade followed by assessment of plasma IFNγ, IL-6, or TNFα by Meso Scale Discovery assay. **A-C**. Pharmacological adrenergic blockade using propranolol (β1/2). **D-F**. Global genetic β1/2 knockout (β1/2^-/-^). Statistics by 2-way ANOVA with Bonferroni post-hoc.

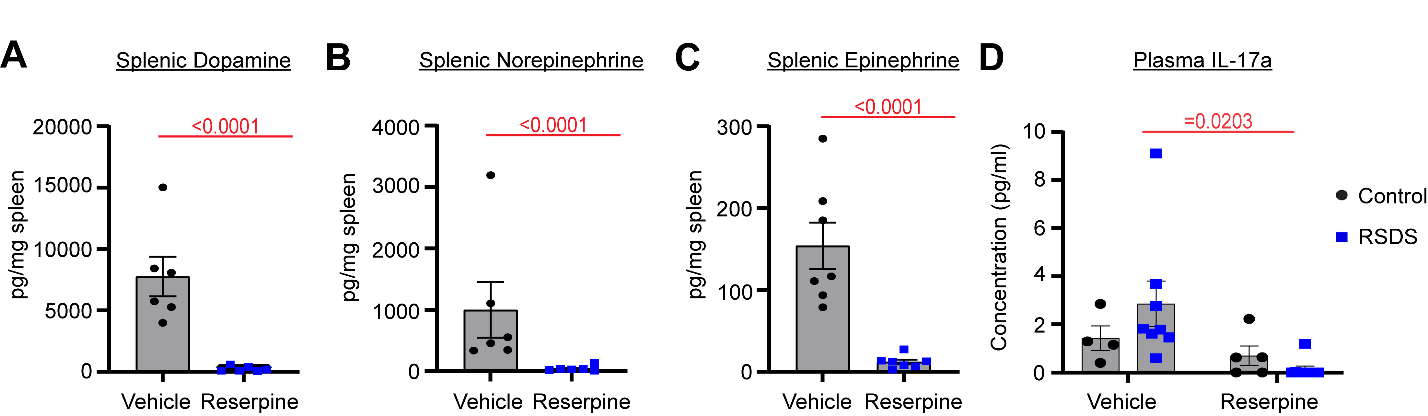

**Supplemental Figure 2. In vivo depletion of catecholamines abrogates IL-17A production.** RSDS was performed in the presence/absence of reserpine followed by assessment of splenic catecholamine levels and IL-17A by Meso Scale Discovery assay. **A-C**. Splenic catecholamines. **D**. Plasma IL-17A. Statistics by Mann-Whitney U test (**A-C**) or 2-way ANOVA with Bonferroni post-hoc (**D**).
